# Moult cycle and setal development of the Atlantic ditch shrimp *Palaemon varians* Leach, 1814 (Decapoda: Caridea: Palaemonidae)

**DOI:** 10.1101/2024.08.23.609361

**Authors:** Kenneth Kim, Jonathan Antcliffe, Allison C. Daley, Marc Robinson-Rechavi

**Affiliations:** Department of Ecology and Evolution, Université de Lausanne, 1015, Lausanne, Switzerland; Swiss Institute of Bioinformatics, 1015, Lausanne, Switzerland; Institute of Earth Sciences (ISTE), Université de Lausanne, 1015, Lausanne, Switzerland

**Keywords:** Arthropoda, Crustacea, ecdysis, exoskeleton, larval development, moulting, uropod, setae

## Abstract

The Atlantic ditch shrimp *Palaemon varians* Leach, 1814 is a common estuarine and brackish water species on Northern Atlantic coasts. *Palaemon varians* is an appealing model organism for studying arthropod developmental processes, such as moulting (ecdysis). Detailed morphological information on its moult cycle is still lacking, hence we have characterised the changes in the setal features corresponding to the moult stages of *P. varians* grown under laboratory conditions. The stages of the moult cycle were differentiated and described using microscopic analysis of the setae in the uropods of *P. varians* based on Drach’s classification system. Moult stages were defined as early and late post-moult (A and B), inter-moult (C), early-, mid- and late pre-moult (D_0_, D_1,_ and D_2_), as well as ecdysis stage (E), the actual shedding of the exuvia. Average moult cycle duration was 8.7 days, where pre-moult accounted for the longest duration of 4.4 days on average. This study provides a morphological reference for determining the moult stage of *P. varians* without the use of invasive techniques, and thus it is well suited for repetitive observations of an individual to track the entire moulting process.

## INTRODUCTION

Arthropods such as crustaceans need to moult periodically in order to grow. Moulting is a complex and tightly regulated event, governed primarily by two group of hormones, the ecdysteroids and sesquiterpenoids (Campli *et al*., 2024). Both hormones play a crucial role in coordinating the various stages of moulting from the preparatory events leading to and after ecdysis (Cheong *et al*., 2015; Qu *et al*., 2018; Truman & Riddiford, 2019; Campli *et al*., 2024). In decapod crustaceans, the progression of each of these stages is initiated and coordinated by ecdysteroids synthesised in the Y-organ, which is negatively regulated by the moult-inhibiting hormone (MIH) and crustacean hyperglycemic hormone (CHH), synthesised in the X-organ/sinus gland (XO-SG) complex in the eyestalk ganglia (Skinner, 1985; Mykles, 2011; Hopkins, 2012; Webster *et al*., 2012). Hemolymph ecdysteroid titers increase during the premoult stage, the old cuticle is degraded, calcium and minerals are reabsorbed, and the new cuticle is synthesized (McCarthy & Skinner, 1977; Skinner, 1985; Skinner *et al*., 1992; Hopkins & Das, 2015). The decrease in hemolymph ecdysteroid titers triggers a stereotyped movement and muscle contraction referred to as the ecdysis behaviour (White & Ewer, 2014), ultimately leading to the exit of the individual from its exuvium through sutural exit gapes (Daley & Drage, 2016). Successful emergence prompts post-ecdysial events such cuticle tanning, mineralization and sclerotization turning the soft and pale covering into a rigid exoskeleton. Given the intricate interplay of hormonal, biochemical, and behavioural processes during moulting, it is crucial to accurately determine the moult stage of an individual in experimental studies. Identifying the moult stage is essential for studying the temporal aspects of hormonal regulation, gene expression patterns, and specific physiological responses related to moulting (Mykles & Chang, 2020).

The moult cycle of Crustacea is divided into the four main stages: post-moult, intermoult, pre-moult, and ecdysis or exuviation, the actual shedding of the old cuticle (Drach, 1939; Spindler *et al*., 1974). Each moult stage can be determined in various ways such as measuring hemolymph ecdysteroid titer (Anger & Spindler, 1987; Snyder & Chang, 1991; Martin-Creuzburg *et al*., 2007; Styrishave *et al*., 2008; Mykles, 2011; Techa & Chung, 2015), gastrolith size (Shechter *et al*., 2008; Gramitto, 1998), and the structural changes of the exoskeleton and various epithelial structures (Drach & Tchernigovtzeff, 1967; review in Gorissen & Sandeman, 2022). Among these, one of the most widely used approaches to determine the moult stage of an animal is by examining the structural changes in its setae. Setogenesis, the formation of new setae, was initially described for moult staging of the brachyuran crab *Cancer pagurus* Linnaeus, 1758 by Drach (1939) and *Palaemon serratus* Pennant, 1777 by Drach & Tchernigovtzeff (1967). Drach’s moult stage classification, which describes the main moulting stages and their corresponding substages, includes the post-moult stage, divided into the early (stage A) and late post-moult (stage B), intermoult (stage C), pre-moult (stage D and substages referred to as D_n_), and ecdysis (stage E). Drach’s staging system has been widely adapted to other crustaceans such as brine shrimps (Criel & Walgraeve, 1989), barnacles (Davis *et al*., 1973), amphipods (Graf, 1986), euphausiids (Buchholz, 1982, 1991; Buchholz & Buchholz, 2010), stomatopods (Reaka 1975), crayfishes (Stevenson, 1968; Mills & Lake, 1975), lobsters (Aiken, 1973; Lyle & Macdonald, 1983; Musgrove, 2000; Marco, 2012), crabs (Sugumar *et al*., 2013) and shrimps (Freeman & Bartell 1975; Robertson *et al*., 1987; Corteel *et al*., 2012; Foguesatto *et al*., 2019).

In order to use Drach’s classification system, it is necessary to first characterise the morphologically recognizable stages of the moult cycle, which may then be used as a reference system for other studies of the species of interest. Challenges can arise when there is a lack of images and detailed descriptions of setal structures for target taxa. *Palaemon varians* Leach, 1814 is one such species for which descriptions of the moult cycle and setogenesis are lacking. Addressing this gap is crucial not only for providing a reference for moult stage determination within the species, but also for contributing valuable data to initiatives such as MoultDB (www.moultdb.org), which aims to compile traits and genomics of moulting across diverse arthropods. Such moult cycle descriptions will ultimately support broader comparative analyses of moulting across crustaceans and other arthropods.

The Atlantic ditch shrimp *P. varians* is a common estuarine and brackish water species found on North Atlantic coasts. The species, which was formerly known as *Palaemonetes varians* (De Grave & Ashelby, 2013) is eurythermal and displays an abbreviated life cycle consisting of at least four larval instars (Oliphant & Thatje, 2021). *Palaemon varians* is an appealing candidate for studying arthropod developmental processes, such as moulting, owing to its well understood ontogenetic series, amenability to breeding in the laboratory, and wide tolerance of aquarium conditions. We determined the moult cycle and the duration of each stage of *P. varians* grown under laboratory conditions and provided specific anatomical description of the changes in its setae for each of the stage and substages using a simple and non-invasive technique.

## MATERIALS AND METHODS

The shrimp used in this study were reared in an aquarium research laboratory of the Institute of Earth Sciences, University of Lausanne. Tanks containing the shrimp were maintained with a water temperature of 21 °C and salinity of 25.1 psu (∼25.1 ppt or a specific gravity of 1.019). We used biofiltration to stabilise a large 300-l tank for the adult population, while regular water changes were required for the small tanks that hosted juveniles. This procedure kept total ammonia-N below 0.5 mg 1^−1^, nitrite-N below 0.15 mg 1^−1^, and nitrates as close to zero as possible. All tanks received artificial sunlight for 8 h per day. Breeding saltwater shrimp in captivity has some challenges and the protocol for the breeding program of this species was optimised as follows. Ovigerous (berried) females were isolated in a nursery tank until the eggs hatched, at which point the adult female was immediately removed and placed back in the main tank. Newly hatched larvae were kept isolated in the nursery tank (12 l) that was aerated using an air pump but had no filtration, as the larvae would otherwise be drawn into the filter. They were fed once daily with *Artemia* nauplii for three weeks and were then weaned onto a commercial pelleted feed (JBL Pronovo, Neuhofen, Germany). Once they grew to 1–1.5 cm in length, they were moved to another small tank (20 l) with filtration, and when they reached 2 cm or greater in length, they were introduced into the main adult-population tank (300 l). The sizes of the shrimp in the tanks were visually estimated in reference to a subset of individuals that were directly measured. Mixing life stages caused cannibalistic predation so they were kept isolated until of appropriate size.

Moult stages of the shrimp were differentiated and characterised by observing the setae of uropods based on Drach & Tchernigovtzeff (1967) and Foguesatto *et al*. (2019). A total of 30 adult shrimp, males and non-ovigerous females, measuring 2.5– 4.5 cm were used for the setal observation. Each shrimp was measured using a ruler from the tip of the rostrum to the tip of the uropods. Ovigerous females were not included in the study since their moult cycle gets arrested during intermoult and they undergo moulting a day after oviposition. During each of the observation periods, shrimp were kept in a separate tank (21°C, 25.1 psu, 20 l, no filtration, air pump used) and were followed individually each day for the duration of at least one entire moult cycle. Following the methodology of Stevenson *et al*. (1968), individuals were observed by wrapping the shrimp in a wet piece of paper cloth or tissue paper with its tail protruding and placed in a Petri plate with water taken from the same tank for observation. Observation time for each shrimp under the microscope was kept to less than 3 min to avoid stressing the shrimp. The setae of uropods were observed using an Olympus SZX10 stereomicroscope using an observed magnification of 0.63–6.3× and were imaged using an SC50 5-megapixel colour 118 camera (Olympus Life Science Solutions, Olympus, Tokyo, Japan) with Preciv Image Analysis Software (version 1.2; Evident Corporation, Tokyo, Japan). The terminology for anatomical structures used to describe setogenesis and moult stages followed that of Foguesatto *et al*. (2019).

## RESULTS

The duration of the moult cycle of *Palaemon varians* ranged 6–12 d, with a mean duration of 8.7 ± 1.4 d (*N* = 30) (Fig. 1), and with smaller individuals moulting faster than larger ones (Pearson correlation coefficient: r = 0.93, *P* < 0.0001). Individuals 2.1–3.1 mm long had a shorter moult duration of 6–8 d, those 3.1–4.0 mm long moulted within 9–10 d, and those longer than 4.0 mm with a moult duration of 10–12 d (Supplementary material Table S1).

**Figure 1.**
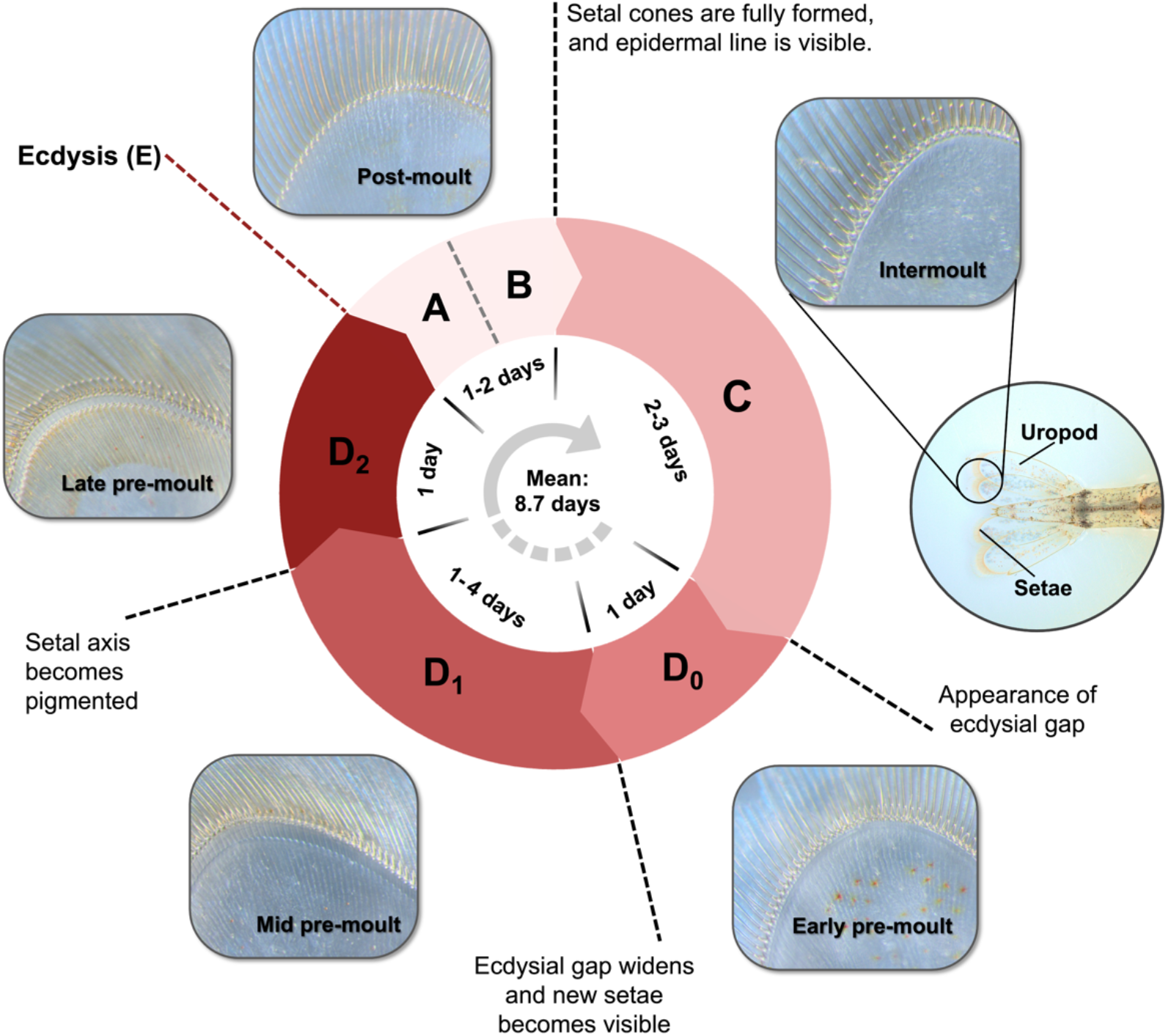
Overview of the moult cycle of *Palaemon varians*, with the duration (in days) of each stage and substage, namely the post-moult stage which is divided into early (Stage A) and late post-moult (Stage B), intermoult (Stage C), pre-moult divided into early (Stage D_0_), mid (Stage D_1_), and late pre-moult (Stage D_2_). Ecdysis stage (E) refers to the exit of the shrimp from its exuvium which takes place 1 to 2 d after late pre-moult (D_2_).

### Post-moult (stages A and B)

After ecdysis, the cuticle of the uropod setae of *P. varians* had a pale appearance with no visible internal features. During early post-moult (stage A), the setae did not have setal cones and the setal bases were transparent and had a slightly rounded appearance. The setal matrix reached the base of the setae with no visible epidermal line (Fig. 2A). Most individuals reached intermoult after 24 h except for the larger ones (> 4 cm), which took up to 48 hours. During late post-moult stage (stage B), the setal bases had a more rounded appearance and the setal matrix still reached the setal base with no visible epidermal line in-between. Setal cones had not fully formed in all of the setae (Fig. 2B). Post-moult stage had a duration of 1–2 d (mean 1.13 ± 0.35 d, *N* = 30).

**Figure 2.**
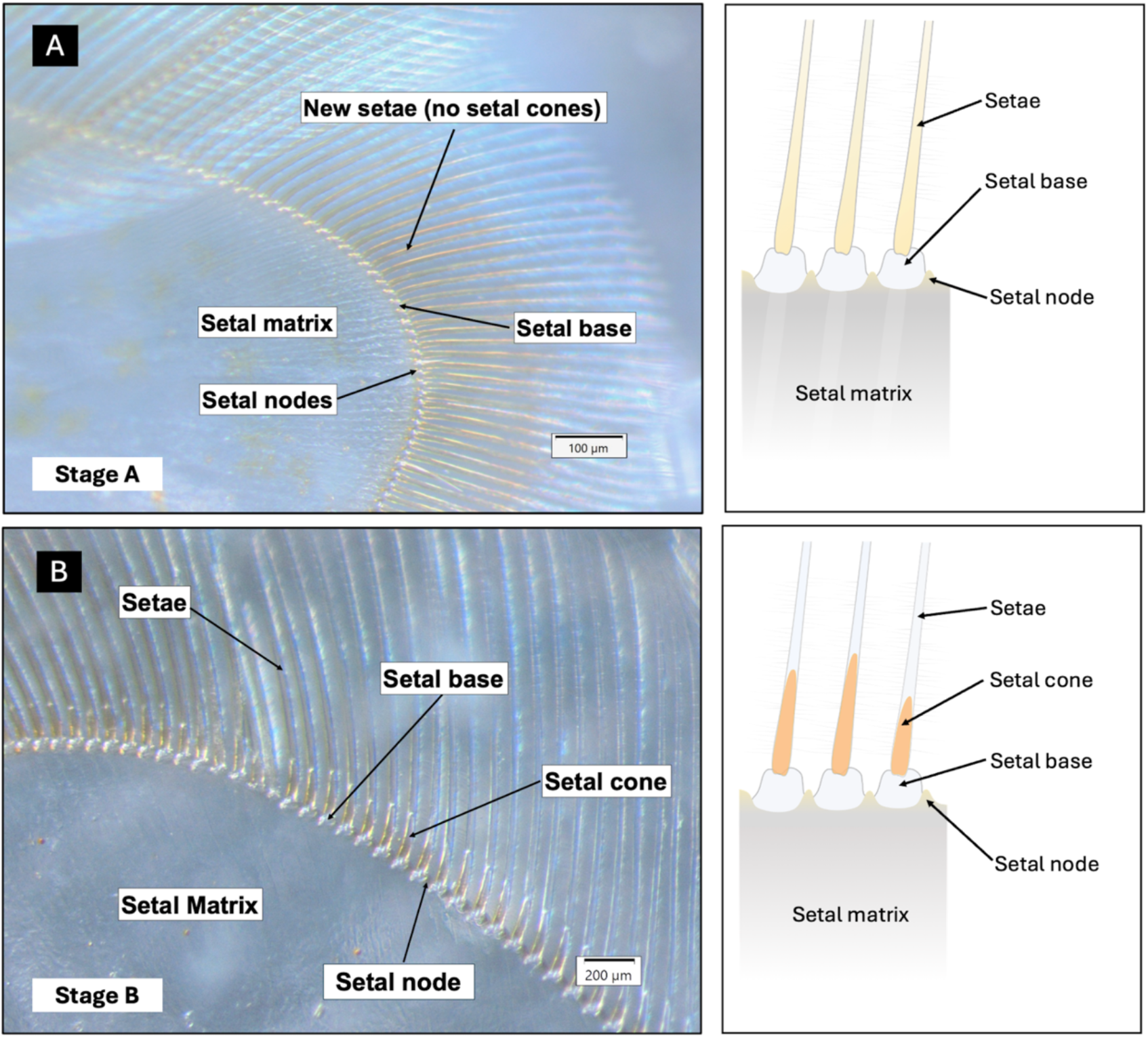
Microscope image (left) and simplified illustration (right) of the uropod setae of *Palaemon varians* during early post-moult (**A**) and late post-moult (**B**).

### Intermoult (stage C)

The setal structures of the uropods were well developed during intermoult. The setal cone was visible at the base of all setae and the epidermal line parallel to the base of the setae was visible. The epidermal line during intermoult closely followed the contour of the base of the setae and no gap was observed. Faint parallel lines could also be observed in the setal matrix where the setal axis eventually formed during the pre-moult stage (Fig. 3). The total duration of the intermoult stage was around 2–3 d (mean 2.8 ± 0.38 d, *N* = 30). This was shorter than the observed intermoult period for *Palaemonetes argentinus* Nobili, 1901, with a mean of 4.8 d (Foguesatto *et al*. 2019) and 4–6 d (Díaz *et al*., 1998).

**Figure 3.**
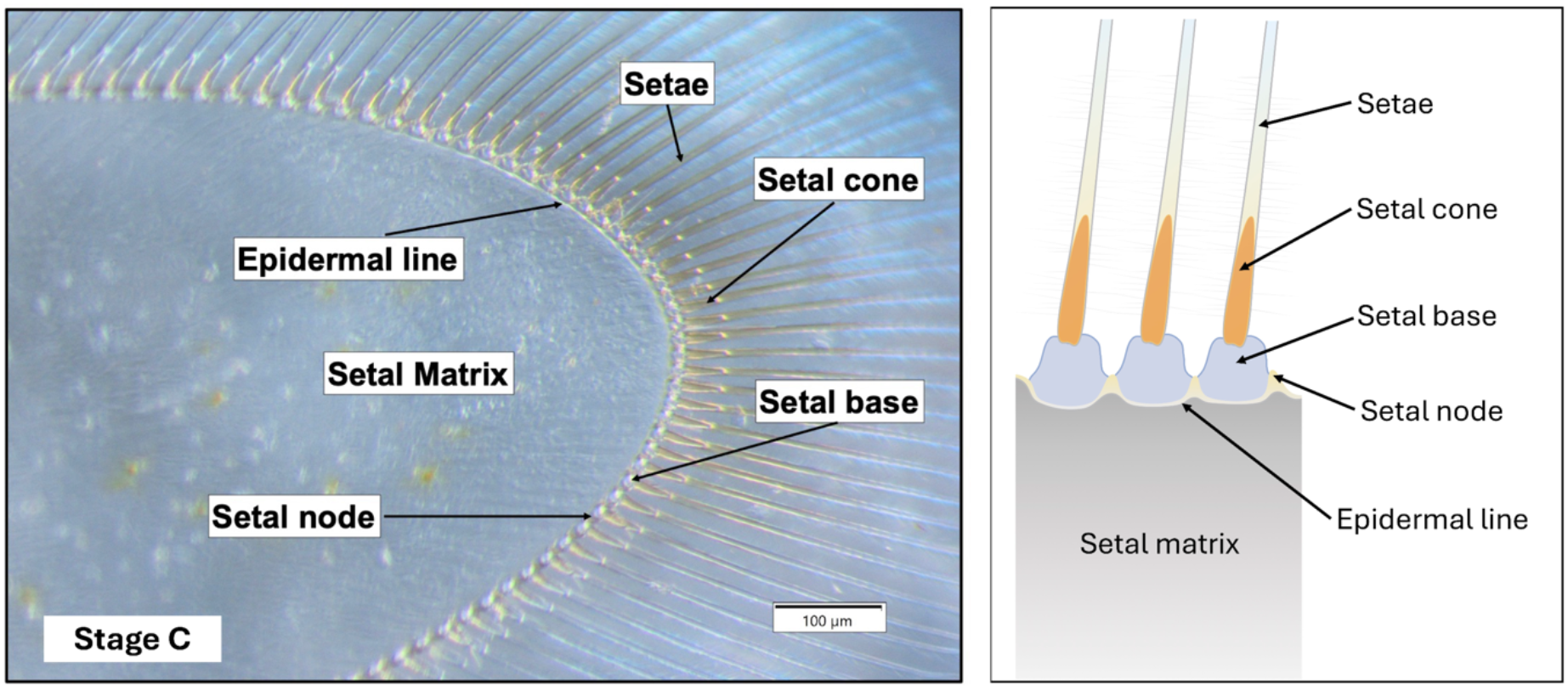
Microscope image (left) and simplified illustration (right) of the uropod setae of *Palaemon varians* during intermoult.

### Pre-moult (stage D)

The early pre-moult phase (D_0_) was marked by the appearance of an ecdysial gap, created by the retraction of the epidermal line away from the setal nodes as the epidermis detached from the old exoskeleton (apolysis) (Fig. 4A). Within the setal matrix, the epidermis around the setal axes was not yet invaginated and the tips of the new setae could already be observed although it only becomes clearly visible in the next stages. The new setae within the ecdysial gap formed by apolysis and the invagination of the epidermis around the setal axes became visible during mid-premoult (D_1_) (Fig. 4B). The progression of mid pre-moult towards the late pre-moult stage was characterised by the invagination of the epidermis around the setal axes, the widening of the ecdysial gap, and with the new setae and the setal axes being more visible. The late pre-moult phase (D_2_) was marked by the pigmentation of the setal axes, which also exhibited a fully formed tube-like appearance (Fig. 4C). The late pre-moult phase usually lasted 24–48 h until ecdysis. The total duration of the pre-moult stage was 3–7 d (mean 4.4 ± 1.09 d, *N* = 30).

**Figure 4.**
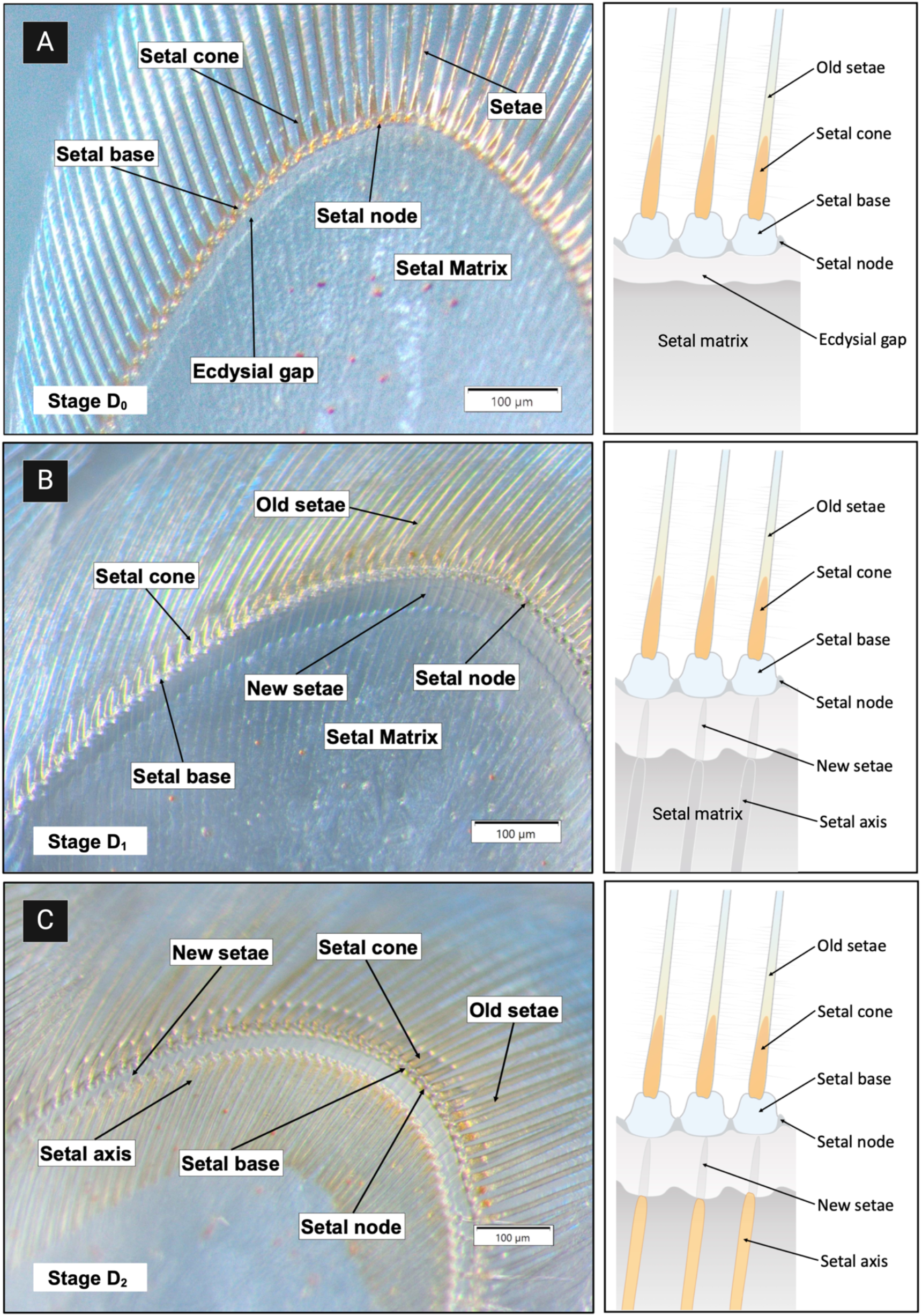
Microscope image (left) and simplified illustration (right) of the uropod setae of *Palaemon varians* during early-, mid- and late pre-moult respectively (**A**–**C**).

### Ecdysis (stage E)

This stage consisted of the act of exuviation, with the shrimp shedding their old exoskeleton. Moulting occurred in just a few seconds to minutes. Moulting and the behaviour immediately prior to and following exuviation was described in *P. varians* by Jefferies (1964), which was confirmed by our observations. Discarded exuviae showed that the ecdysial suture opened between the carapace of the cephalothorax and the abdomen along the dorsal and lateral margins. The cephalothorax and the antennae were gradually withdrawn with an upward (dorsal) motion, after which the posterior body left the exuviae with a rapid flick of the abdomen.

## DISCUSSION

The observed moult cycle duration of *Palaemon varians* closely agrees with values reported for several penaeid shrimps, including 11.5 d in *Penaeus setiferus* (Linnaeus, 1767), 13.6 d in *Penaeus stylirostris* Stimpson, 1874 (Robertson *et al*., 1987), 12.3 d in *Penaeus monodon* Fabricius, 1798 and 10.9 d in *Penaeus vannamei* Boone, 1931 (Corteel *et al*., 2012). The cycle is relatively shorter than the 19 d reported for *Palaemonetes argentinus* (Foguesatto *et al*., 2019). In addition to size differences, several extrinsic factors such as temperature, salinity, nutrition, and water chemistry may influence moulting duration (Lemos & Weissman, 2021). Even within the same species, moulting duration thus exhibited differences in separate studies, as observed in *Palaemonetes argentinus* (Díaz *et al*., 1998; Foguesatto *et al*., 2019) and *Penaeus* m*onodon* (Chan *et al*., 1988; Corteel *et al*., 2012). Even though the relative duration of the moult cycle of *P. varians* is shorter than in *P. argentinus*, the defining structural changes of its setae during the moulting process are almost the same. No species nor sex-specific structural changes were observed during the moult cycle stages between *P. setiferus* and *P. stylirostris* except for the duration of the moult stages (Robertson *et al*., 1987).

The formation of a gap caused by the retraction of the epidermis from the cuticle was observed during early premoult (stage D_0_). This is the main defining characteristic of the beginning of premoult as in other shrimp species (Robertson *et al*., 1987; Chan *et al*., 1988; Díaz *et al*., 1998; Corteel *et al*., 2012; Foguesatto *et al*., 2019). The duration of the premoult stage, however, is longer for the other shrimp species, making it much easier to detect and to assign specific substages. D_0_ is characterised only by the formation of apolysis in *P. argentinus* (Foguesatto *et al*., 2019). The emergence of the new setae could already be seen but it was not yet as evident as in the next stages, which is similar in *P. setiferus* and *P. stylirostris*. Instead of having a separate D_0_ and D_1_, the moult stage was therefore referred to as D_0_ to D_1_, both categorised as early premoult (Robertson *et al*., 1987). Despite having overlapping D_0_ and D_1_, the D_1_ stage in *P. varians* exhibited more pronounced characteristics, including a wider ecdysial gap, the invagination of the epidermis around the setal axis, and the new setae being more protruded and visible. The later observation is similar to the progression of D_0_ to D_1_ observed in *Palaemon pugio* (Holthuis, 1949), where the retraction of the epidermis widens toward late D_0_, and the invaginations further develop in D_1_ (Freeman & Bartell, 1974). In contrast to *P. varians*, where the emergence of the new setae already starts towards the late D_0_, this development is only observed towards the end of D_1_ in *P. pugio* and is only visible in the last stage of premoult (D_2_). The defining characteristic that is observed in *P. varians* during D_2_ is the pigmentation of the setal axes, with the setal axes exhibiting a tubular appearance similar to *P. argentinus*. By contrast, no pigmentation of the setal axes or setal-forming regions in D_2_ was reported in *P. pugio* by Freeman & Bartell (1974). No further changes were observed during stage D_2_ until ecdysis that would justify adding stages D_3_ or D_4_, as observed in other shrimp species (Chan *et al*., 1988; Hunter & Uglow, 1998; Foguesatto *et al*., 2019). The main defining feature of *P. varians* during the post-moult stage is the absence of setal cones, which is similar to other shrimp species (Robertson., *et al* 1987; Chan *et al*., 1988; Díaz *et al*., 1998; Corteel *et al*., 2012; Foguesatto *et al*., 2019). The vesicular inclusions characteristic of the post-moult stage in *P. argentinus*, however, were not observed in *P. varians*. No definitive criteria were observed between early and late post-moult except for the formation of setal cones.

We determined the moult cycle of *P. varians* without the use of invasive techniques. such as cutting off appendages, since these are known to influence moulting rate in crustaceans (Skinner, 1985). We were able to characterise early and late post-moult (stages A and B), inter-moult (stage C), and early, mid and late pre-moult (stages D_0_, D_1_, and D_2_) using the setal changes in the uropods. Discrepancies and differences in the moult stage characteristics of *P. varians* in comparison to other shrimp species and across Decapoda are observed, such as during the mid pre-moult and transition of late post-moult to intermoult. Our findings are consistent with the general trend of moult staging criteria amongst decapods, in which intermoult (stage C) is characterised by the presence of setal cones and the absence of apolysis, early premoult (D_0_) is characterised by formation of apolysis, mid-premoult (D_1_) by setal invagination and pigmentation of the setal axis in late pre-moult (D_2_). While there are no consistent criteria to define early and late post-moult, the absence of setal cones appear to be widely observed amongst shrimp and crayfish species (Gorissen & Sandeman, 2022).

Our study, along with previous work on decapods, has provided a comprehensive morphological reference and nomenclature for moult stage classification (reviewed in Gorissen & Sandeman, 2022). While many of these criteria have been widely used to compare moult stages across Decapoda, certain traits, such as the formation of the ecdysial gap, are products of conserved processes like apolysis, observed not only in crustaceans but across Arthropoda as a whole. Addressing how comparable are these moult criteria amongst crustaceans will be essential for broader comparative studies of moulting across Arthropoda.

## ACKNOWLEDGEMENTS

The authors would like to thank the members of the Arthropod Moulting Evolution Project: Robert Waterhouse, Giulia Campli, Michele Leone, Sagane Joye-Dind, Valentine Rech de Laval, Ariel Chipman, Olga Volovych, Idan Sheizaf, Asya Novikova, Sinead Lynch, and Harriet Drage, for their invaluable support, assistance, and insightful discussions throughout this study. We are grateful to Christian de Guttry, Claudia Baumgartner, Farid Saleh, and Nora Corthésy for their help in the aquarium laboratory and with equipment use. Finally, we extend our gratitude to the anonymous reviewers for their constructive feedback and thoughtful suggestions, which greatly improved the manuscript. This work was supported by funding from a Swiss National Science Foundation (SNSF) Sinergia award [grant number 198691] and an SNSF Careers award [grant number 202669]. Figures were created with Biorender.com under Biorender’s academic license terms.

## SUPPLEMENTARY MATERIAL

Supplementary material is available at *Journal of Crustacean Biology* online.

S1 Table. Duration in days of each of the moult stage of *Palaemon varians*.

S2 Figure. Discarded exuvium of *Palaemon varians*.

## REFERENCES

Aiken, D.E. 1973. Proecdysis, setal development, and molt prediction in the American lobster (Homarus americanus). Journal of the Fisheries Research Board of Canada, 30: 1337–1344.

Anger, K. & Spindler, K.D. 1987. Energetics, moult cycle and ecdysteroid titers in spider crab (Hyas araneus) larvae starved after the D_0_threshold. Marine Biology, 94: 367–375.

Boone, L. 1931. A collection of anomuran and macruran Crustacea from the Bay of Panama and the fresh waters of the Canal Zone. Bulletin of the American Museum of Natural History, 63: 137–189.

Buchholz, F. 1982. Drach’s molt staging system adapted for euphausiids. Marine Biology, 66: 301–305.

Buchholz, F. 1991. Moult cycle and growth of Antarctic krill (Euphausia superba) in the laboratory. Marine Ecology Progress Series, 69: 217–229.

Buchholz, F. & Buchholz, C. 2010. Growth and moulting in northern krill (Meganyctiphanes norvegica Sars). Advances in Marine Biology, 57: 173– 197.

Campli, G., Volovych, O., Kim, K., Veldsman, W.P., Drage, H.B., Sheizaf, I., Lynch, S., Chipman, A.D., Daley, A.C., Robinson-Rechavi, M. & Waterhouse, R.M. 2024. The moulting arthropod: a complete genetic toolkit review. Biological Reviews. [10.1111/brv.13123].

Chan, S.M., Rankin, S.M. & Keeley, L.L. 1988. Characterization of the molt stages in Penaeus vannamei: setogenesis and hemolymph levels of total protein, ecdysteroids, and glucose. Biological Bulletin, 175: 185–192.

Cheong, S.P.S., Huang, J., Bendena, W.G., Tobe, S.S. & Hui, J.H.L. 2015. Evolution of ecdysis and metamorphosis in arthropods: The rise of regulation of juvenile hormone. Integrative and Comparative Biology, 55: 878–890.

Corteel, M., Dantas-Lima, J.J., Wille, M., Alday-Sanz, V., Pensaert, M.B., Sorgeloos, P. & Nauwynck, H.J. 2012. Moult cycle of laboratory-raised Penaeus (Litopenaeus) vannamei and P. monodon. Aquaculture International, 20: 13–18.

Criel, G.R.J. & Walgraeve, H.R.M.A. 1989. Molt staging in Artemia adapted to Drach’s system. Journal of Morphology, 199: 41–52.

Daley, A.C. & Drage, H.B. 2016. The fossil record of ecdysis, and trends in the moulting behaviour of trilobites. Arthropod Structure and Development, 45: 71–96.

Davis, C.W., Fyhn, U.E.H. & Fyhn, H.J. 1973. The intermolt cycle of cirripeds: Criteria for its stages and its duration in Balanus amphitrite. Biological Bulletin, 145: 310–322.

De Grave, S. & Ashelby, C.W. 2013. A re-appraisal of the systematic status of selected genera in Palaemoninae (Crustacea: Decapoda: Palaemonidae). Zootaxa, 3734: 331–344.

Díaz, A., Petriella, A.M. & Sousa, L. 1998. Setogenesis and growth of the freshwater prawn Palaemonetes argentinus (Decapoda, Caridea, Palaemonidae). Iheringia Série Zoologia, 85: 59–65.

Drach, P. 1939. Mue et cycle d’intermue chez les Crustacés Décapodes. Annales de l’Institut Océanographique, 19: 103–391.

Drach, P. & Tchernigovtzeff, C. 1967. Sur la méthode de détermination des stades d’intermue et son application générale aux crustacés. Vie et Milieu, 18: 595–610.

Fabricius, J.C. 1798. Supplementum Entomologiae Systematicae. Proft & Storch, Hafniae [= Copenhagen].

Foguesatto, K., Nery, L.E.M. & Souza, M.M. 2019. Setogenesis and characterization of the new moult substages in the freshwater shrimp Palaemon argentinus (Nobili, 1901) (Caridea: Palaemonidae). Nauplius, 27: e2019013.[10.1590/2358-2936e2019013].

Freeman, J.A. & Bartell, C.K. 1975. Characterization of the molt cycle and its hormonal control in Palaemonetes pugio (Decapoda, Caridea). General and Comparative Endocrinology, 25: 517–528.

Gorissen, S. & Sandeman, D. 2022. Moult cycle staging in decapod crustaceans (Pleocyemata) and the Australian crayfish, Cherax destructor Clark, 1936 (Decapoda, Parastacidae). Crustaceana, 95: 165–217.

Graf, F. 1986. Fine determination of the molt cycle stages in Orchestia cavimana Heller (Crustacea: Amphipoda). Journal of Crustacean Biology, 6: 666–678.

Gramitto, M.E. 1998. Molt pattern identification through gastrolith examination on Nephrops norvegicus (L.) in the Mediterranean Sea. Scientia Marina, 62: 17–23.

Holthuis, L.B. 1949. Note on the species of Palaemonetes (Crustacea Decapoda) found in the United States of America. Proceedings of the Koninklijke Nederlandse Akademie van Wetenschappen, 52: 87–95.

Hopkins, P.M. 2012. The eyes have it: A brief history of crustacean neuroendocrinology. General and Comparative Endocrinology, 175: 357– 366.

Hopkins, P.M. & Das, S. 2015. Regeneration in crustaceans, pp. 168–198. In: The Natural History of the Crustacea, Vol. 4 (E.S. Chang & M. Thiel, eds.). Oxford University Press, Oxford.

Hunter, D.A. & Uglow, R.F. 1998. Setal development and moult staging in the shrimp Crangon crangon (L.) (Crustacea: Decapoda: Crangonidae). Ophelia, 49: 195–209.

Jefferies, D.J. 1964. The moulting behaviour of Palaemonetes varians (Leach) (Decapoda; Palaemonidae). Hydrobiologia, 24: 457–488.

Leach, W.E. 1814. Crustaceology. In: Brewster’s Edinburgh Encyclopaedia, Vol. 7 (D. Webster, ed.). Balfour, Edinburgh.

Lemos, D. & Weissman, D. 2021. Moulting in the grow-out of farmed shrimp: a review. Reviews in Aquaculture, 13: 5–17.

Linnaeus, C. 1758. Systema Naturae per Regna Tria Naturae, Secundum Classes, Ordines, Genera, Species, cum Characteribus, Differentiis, Synonymis, Locis, Vol. 1, Edn. 10. Reformata. Laurentii Salvii, Holmiae [= Stockholm].

Linnaeus, C. 1767. Systema Naturæ per Regna tria Naturæ, secundum Classes, ordines, Genera, Species, cum Characteribus, Differentiis, Synonymis, Locis, Vol. 1, Edn. 13. Vindobona [=Vienna].

Lyle, W.G. & MacDonald, C.D. 1983. Molt stage determination in the Hawaiian spiny lobster Panulirus marginatus. Journal of Crustacean Biology, 3: 208–216.

Marco, H.G. 2012. The moult cycle of Jasus lalandii (H. Milne Edwards, 1837) (Decapoda, Palinuridae), a cold water spiny lobster species: changes in ecdysteroid titre and setogenesis. Crustaceana, 85: 1221–1239.

Martin-Creuzburg, D., Westerlund, S.A. & Hoffmann, K.H. 2007. Ecdysteroid levels in Daphnia magna during a molt cycle: Determination by radioimmunoassay (RIA) and liquid chromatography–mass spectrometry (LC–MS). General and Comparative Endocrinology, 151: 66–71.

McCarthy, J.F. & Skinner, D.M. 1977. Interruption of proecdysis by autotomy of partially regenerated limbs in the land crab, Gecarcinus lateralis. Developmental Biology, 61: 299–310.

Mills, B. & Lake, P. 1975. Setal development and moult staging in the crayfish Parastacoides tasmanicus (Erichson) (Decapoda: Parastacidae). Marine and Freshwater Research, 26: 103–107.

Musgrove, R.J.B. 2000. Molt staging in the southern rock lobster Jasus edwardsii. Journal of Crustacean Biology, 20: 44–53.

Mykles, D.L. 2011. Ecdysteroid metabolism in crustaceans. Journal of Steroid Biochemistry and Molecular Biology, 127: 196–203.

Mykles, D.L. & Chang, E.S. 2020. Hormonal control of the crustacean molting gland: Insights from transcriptomics and proteomics. General and Comparative Endocrinology, 294: 113493 [10.1016/j.ygcen.2020.113493].

Nobili, G., 1901. Decapodi raccolti dal Dr. Filippo Silvestri nell’America meridionale. Bolletino dei Musei di Zoologia ed Anatomia comparata della R. Università di Torino, 16: 1–16.

Oliphant, A. & Thatje, S. 2021. Variable shrimp in variable environments: reproductive investment within Palaemon varians. Hydrobiologia, 848: 469–484.

Pennant, T. 1777. British Zoology, Vol. 4, Crustacea.Mollusca. Testacea. B. White, London.

Qu, Z., Bendena, W.G., Tobe, S.S. & Hui, J.H.L. 2018. Juvenile hormone and sesquiterpenoids in arthropods: biosynthesis, signaling, and role of MicroRNA. Journal of Steroid Biochemistry and Molecular Biology. 184: 69–76.

Reaka, M.L. 1975. Molting in stomatopod crustaceans. I. Stages of the molt cycle, setagenesis, and morphology. Journal of Morphology, 146: 55–80.

Robertson, L., Bray, W., Leung-Trujillo, J. & Lawrence, A. 1987. Practical molt staging of Penaeus setiferus and Penaeus stylirostris. Journal of the World Aquaculture Society, 18: 180–185.

Shechter, A., Berman, A., Singer, A., Freiman, A., Grinstein, M., Erez, J., Aflalo, E.D. & Sagi, A. 2008. Reciprocal changes in calcification of the gastrolith and cuticle during the molt cycle of the red claw crayfish Cherax quadricarinatus. Biological Bulletin, 214: 122–134.

Skinner, D.M. 1985. Moulting and regeneration. pp. 44–146. In: Bliss, D.E. & Mantel, L.H. (eds.), The Biology of Crustacea, Vol. 9. Academic Press, New York.

Skinner, D.M., Kumari, S.S. & O’Brien, J.J. 1992. Proteins of the crustacean exoskeleton. American Zoologist, 32: 470–484.

Snyder, M.J. & Chang, E.S. 1991. Ecdysteroids in relation to the molt cycle of the American lobster, Homarus americanus. General and Comparative Endocrinology, 81: 133–145.

Spindler, K.D., Adelung, D. & Tchernigovtzeff, C. 1974. A comparison of the methods of molt staging according to Drach and to Adelung in the common shore crab, Carcinus maenas. Zeitschrift für Naturforschung C, 29: 754– 756.

Stevenson, J.R. 1968. Metecdysial molt staging and changes in the cuticle in the crayfish Orconectes sanborni (Faxon). Crustaceana, 14: 169–177.

Stevenson, J.R., Guckert, R.H. & Cohen, J.D. 1968. Lack of correlation of some proecdysial growth and developmental processes in the crayfish. Biological Bulletin, 134: 160–175.

Stimpson, W.S. 1874. Notes on North American Crustacea in the Museum of the Smithsonian Institution, No. III. Annals of the Lyceum of Natural History of New York, 10: 92–136.

Styrishave, B., Lund, T. & Andersen, O. 2008. Ecdysteroids in female shore crabs Carcinus maenas during the moulting cycle and oocyte development. Journal of the Marine Biological Association of the United Kingdom, 88: 575–581.

Sugumar, V., Vijayalakshmi, G. & Saranya, K. 2013. Molt cycle-related changes and effect of short-term starvation on the biochemical constituents of the blue swimmer crab Portunus pelagicus. Saudi Journal of Biological Sciences, 20: 93–103.

Techa, S. & Chung, J.S. 2015. Ecdysteroids regulate the levels of molt-inhibiting hormone (MIH) expression in the blue crab, Callinectes sapidus. PLoS ONE, 10: e0117278.[10.1371/journal.pone.0117278].

Truman, J.W. & Riddiford, L.M. 2019. The evolution of insect metamorphosis: a developmental and endocrine view. Philosophical Transactions of the Royal Society B: Biological Sciences, 374: 20190070 [10.1098/rstb.2019.0070].

Webster, S.G., Keller, R. & Dircksen, H. 2012. The CHH-superfamily of multifunctional peptide hormones controlling crustacean metabolism, osmoregulation, moulting, and reproduction. General and Comparative Endocrinology, 175: 217–233.

White, B.H. & Ewer, J. 2014. Neural and hormonal control of postecdysial behaviors in insects. Annual Review of Entomology, 59: 363–381.

