## Supplementary File for "Moult cycle and setal development of the Atlantic ditch shrimp *Palaemon varians* Leach, 1814 (Decapoda: Caridea: Palaemonidae)" (from another version of this preprint: version 1, 2024-08-23)

**Supplementary Table 1.** Counts in days for each moult stage and substages of *Palaemon varians*

| sample_id | Intermoult | Early_premoult | Mid_premoult | Late_premoult | Postmoult | Total | size |
| --- | --- | --- | --- | --- | --- | --- | --- |
| PV_5 | 3 | 1 | 4 | 2 | 2 | 12 | 4.5 |
| PV_12 | 3 | 2 | 3 | 2 | 2 | 12 | 4.1 |
| PV_30 | 3 | 1 | 2 | 2 | 2 | 10 | 4 |
| PV_1 | 3 | 1 | 2 | 2 | 1 | 9 | 3.1 |
| PV_3 | 3 | 1 | 2 | 2 | 1 | 9 | 3.1 |
| PV_8 | 3 | 1 | 2 | 2 | 1 | 9 | 3.1 |
| PV_9 | 3 | 1 | 3 | 1 | 1 | 9 | 3.1 |
| PV_11 | 3 | 1 | 2 | 2 | 1 | 9 | 3.1 |
| PV_15 | 3 | 1 | 2 | 1 | 2 | 9 | 3.1 |
| PV_16 | 3 | 1 | 2 | 2 | 1 | 9 | 3.1 |
| PV_27 | 3 | 1 | 3 | 1 | 1 | 9 | 3.1 |
| PV_28 | 3 | 1 | 2 | 2 | 1 | 9 | 3.1 |
| PV_2 | 3 | 1 | 2 | 1 | 1 | 8 | 3 |
| PV_4 | 3 | 1 | 2 | 1 | 1 | 8 | 3 |
| PV_6 | 3 | 1 | 2 | 1 | 1 | 8 | 3 |
| PV_7 | 3 | 1 | 2 | 1 | 1 | 8 | 3.1 |
| PV_10 | 2 | 1 | 2 | 2 | 1 | 8 | 3 |
| PV_13 | 3 | 1 | 2 | 1 | 1 | 8 | 3.1 |
| PV_14 | 3 | 1 | 2 | 1 | 1 | 8 | 3.1 |
| PV_21 | 3 | 1 | 2 | 1 | 1 | 8 | 3 |
| PV_23 | 3 | 1 | 2 | 1 | 1 | 8 | 3 |
| PV_24 | 3 | 1 | 2 | 1 | 1 | 8 | 3 |
| PV_26 | 3 | 1 | 2 | 1 | 1 | 8 | 3 |
| PV_18 | 3 | 1 | 1 | 1 | 1 | 7 | 2.5 |
| PV_19 | 2 | 1 | 2 | 1 | 1 | 7 | 2.1 |
| PV_20 | 2 | 1 | 2 | 1 | 1 | 7 | 2.5 |
| PV_22 | 3 | 1 | 1 | 1 | 1 | 7 | 2.5 |
| PV_25 | 2 | 1 | 2 | 1 | 1 | 7 | 2.1 |
| PV_29 | 3 | 1 | 1 | 1 | 1 | 7 | 2.1 |
| PV_17 | 2 | 1 | 1 | 1 | 1 | 6 | 2.1 |

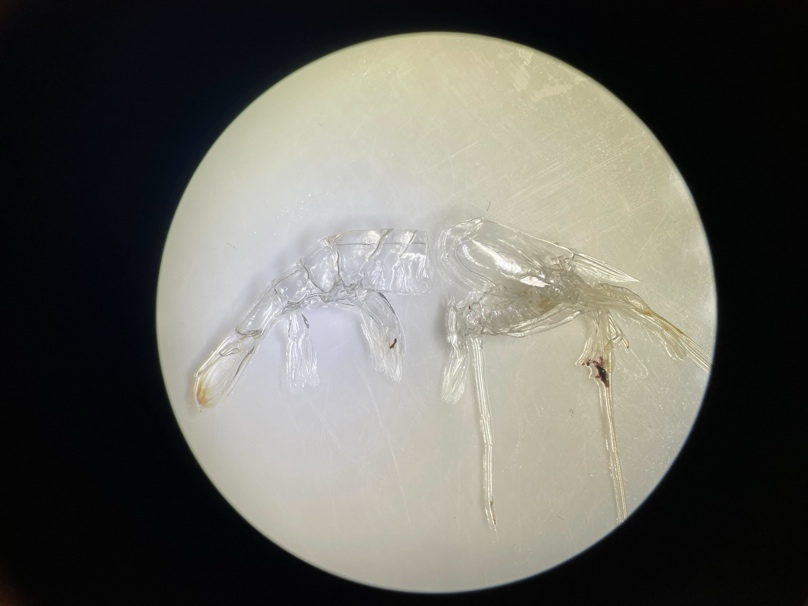

**Supplementary Figure 1.** Discarded exuvia of *Palaemon varians*
